# STAT3 but not STAT5 contributes to the protective effect of electro-acupuncture against myocardial ischemia/reperfusion injury

**DOI:** 10.1101/2020.07.22.215384

**Authors:** Hui-Hui Guo, Xin-Yue Jing, Hui Chen, Hou-Xi Xu, Bing-Mei Zhu

**Author notes:** These authors have equal contribution (Hui-Hui Guo, Xin-Yue Jing). Corresponding author Prof. Bing-Mei Zhu, Present address: Regenerative Medicine Research Center, West China Hospital, Sichuan University, Keyuan Road 4, Gaopeng Street, Chengdu, Sichuan, P.R.China. 610041, (O).

## Abstract

Late remote ischemia preconditioning (RIPC) and electro-acupuncture (EA) have both been suggested to reduce injury caused by myocardial ischemia/reperfusion (I/R). Our previous study has found that cardioprotection in RIPC is STAT5-dependent. Here, we aim to observe the effects of electro-acupuncture pretreatment (EAP) on I/R in the presence or absence of STAT5 in mice and investigate whether the protection of EAP is in a STAT5-dependent manner. In this study, EAP decreased myocardial infarction size (IS) /total area (TA) and rate of cardiomyocyte apoptosis. STAT5 was activated by EAP in the *Stat5*^*fl/fl*^ mice but not in the *Stat5-cKO* mice, whereas, STAT3 was activated by EAP only in the *Stat5-cKO* but not in the *Stat5*^*fl/fl*^ mice. Differentially expressed genes (DEGs) regulated by EAP in the *Stat5*^*fl/fl*^ and the *Stat5-cKO* mice were quite distinct, indicating that EAP may activate IL-6/STAT3 signal in the absence of *Stat5*, and that EAP-induced cardioprotection against myocardial I/R injury was correlated with the activation of anti-apoptotic signaling and cardiomyocyte-survival signaling. Our results, for the first time, demonstrated that the protective effect of EAP was attributed to, but not dependent on, STAT5.

## Introduction

It is known that myocardial ischemia/reperfusion (I/R) injury is a pathological phenomenon that can cause further cardiomyocyte death following blood restoration (Li *et al*, 2019; Chen *et al*, 2019). Remote ischemic preconditioning (RIPC) has been demonstrated as an intervention to attenuate ischemic-reperfusion (I/R) injury and protect myocardium for many years (Davidson *et al*, 2019; Ekeloef *et al*, 2019; Cho & Kim, 2019; Heusch, 2015). Our recent study has shown that signal transducer and activator of transcription 5 (STAT5) plays a critical role in RIPC, and RIPC mediates cardioprotection by activating anti-apoptotic and cardiomyocyte-survival signaling in a STAT5-dependent manner (Chen *et al*, 2018). Electro-acupuncture (EA), as one of the Traditional Chinese Medicine (TCM) approaches, has been applied to treat many diseases clinically around the world (Painovich & Longhurst, 2015). Increasing experimental and clinical evidences have confirmed EA’s effectiveness on cardiovascular diseases (Zhang *et al*, 2020; Ji *et al*, 2018; Huang *et al*, 2014; Fu *et al*, 2014; Zeng *et al*, 2018; Lu *et al*, 2016). A very recent article (Zhao *et al*, 2019) collected a total of 1651 patients with chronic stable angina and performed a clinical trial with acupuncture treatment. This study has shown that acupuncture as an auxiliary treatment method can alleviate clinical pain in patients with chronic stable angina, reduce the patient’s anxiety and depression, and eventually improve the quality of life for these patients. Its underlying mechanisms have been known, include anti-apoptosis and anti-oxidative stress, as well as reducing inflammatory damage, calcium overload, and endoplasmic reticulum stress (Chen *et al*, 2019; Tang *et al*, 2019).

Studies, including our previous work, have shown that using electro-acupuncture before the event of ischemia-reperfusion, which is also called electro-acupuncture pretreatment (EAP), could protect cardiomyocytes by reducing the myocardial infarct size and regulating some molecular signaling, such as apoptosis and survival signaling (Huang *et al*, 2014; Lu *et al*, 2016). Given that the main molecule involved in RIPC in human patients is STAT5 (Cheung *et al*, 2006; Chen *et al*, 2018), we question whether EAP can also protect myocardium against ischemia-reperfusion through STAT5 signaling pathway. Therefore, we have employed the cardiomyocyte-specific *Stat5* knockout (*Stat5-cKO*) mice and generated myocardial I/R model. EA is applied to the mice seven days before the I/R surgery. This study will determine whether EAP shares the same mechanism with RIPC and plays a preconditioning-like role as RIPC does on I/R injury. We have also carried on a genome-wide gene profiling to further find more candidate genes involved in the cardioprotection that results from EAP.

## Results

### 1. EAP reduced myocardial infarct size and attenuated cardiomyocytic apoptosis to the same extent in both the *Stat5* ^*fl/fl*^ mice and the *Stat5-cKO* mice

We did not observe any difference in the daily behavior and cardiac performance between the *Stat5* ^*fl/fl*^ mice and the *Stat5-cKO* mice, same as seen in our previous study (Chen *et al*, 2018). EA pretreatment was applied on PC6 acupoints in both *Stat5* ^*fl/fl*^ mice and *Stat5-cKO* mice for seven days before they received myocardial I/R surgery. We did not see any difference before or after EAP in their daily behavior and cardiac performance between these two genotypes. We then harvested the heart tissues after I/R and measured myocardial infarct areas by TTC staining (Fig 1) and found that EAP significantly reduced infarct size in the *Stat5*^*fl/fl*^ mice (44.6 ± 1.3% without EAP vs. 31.2 ± 5.1% with EAP, *P* < 0.05) and the *Stat5-cKO* mice (41.9 ± 4.1% without EAP vs. 31.3 ± 4.5% with EAP, *P* < 0.05). There was no significant difference between the *Stat5*^*fl/fl*^+EA+I/R and the *Stat5-cKO*+EA+I/R mice.

**Figure 1.**
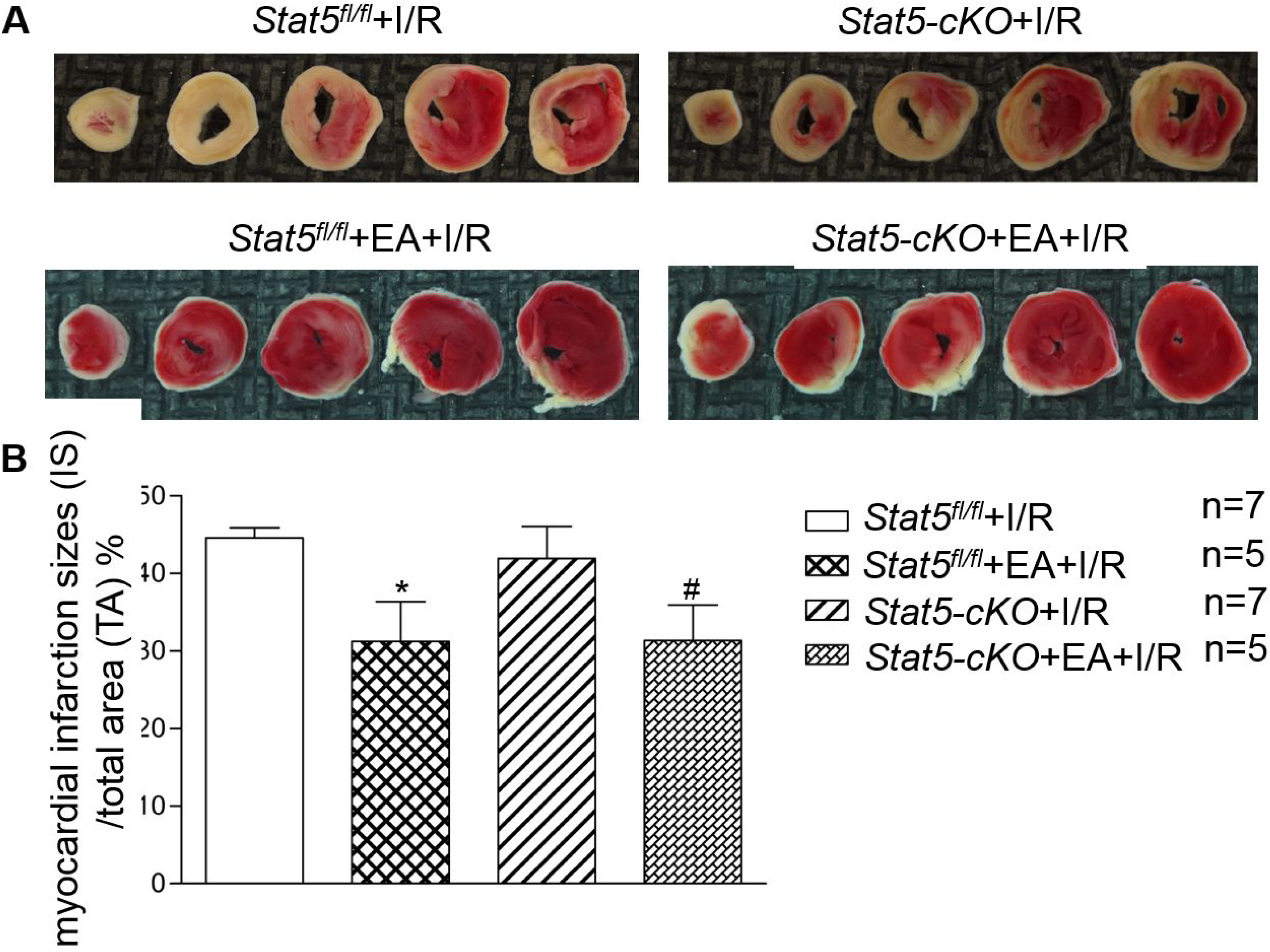
Acupuncture reduced myocardial infarct size. A TTC staining was used to measure ischemic infarct area. B The ratio of infarct size/ total area was calculated and presented as percentage. Data information: In B, data are presented as mean ± SEM, data were analyzed by one-way ANOVA with Tukey’s post hoc correction. Normal tissues are red, and ischemic infarct areas are pale white. \**P* < 0.05, compared with *Stat5*^*fl/fl*^+I/R group, #*P* < 0.05, compared with *Stat5-cKO*+I/R group. Source data are available online for this figure.

TUNEL staining was performed to detect apoptosis in the myocardial cells. As shown in Fig 2, the *Stat5*^*fl/fl*^+EA+I/R group had fewer TUNEL positive cells compared to the *Stat5*^*fl/fl*^+I/R group (*P* < 0.01). Likewise, the apoptotic myocardial cells were also significantly reduced in the *Stat5-cKO*+EA+I/R group compared to the *Stat5-cKO*+I/R group (*P* < 0.01).

**Figure 2.**
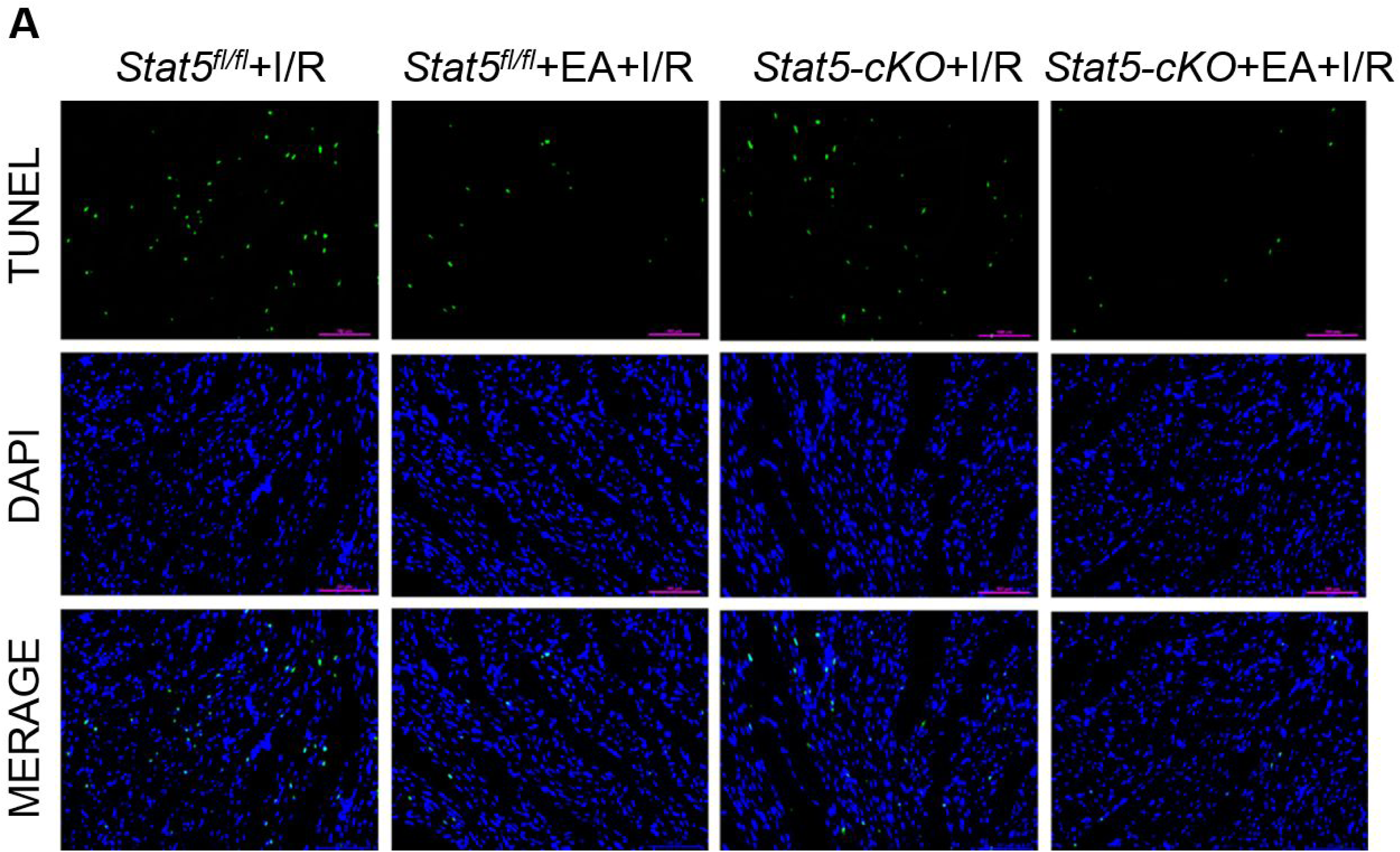
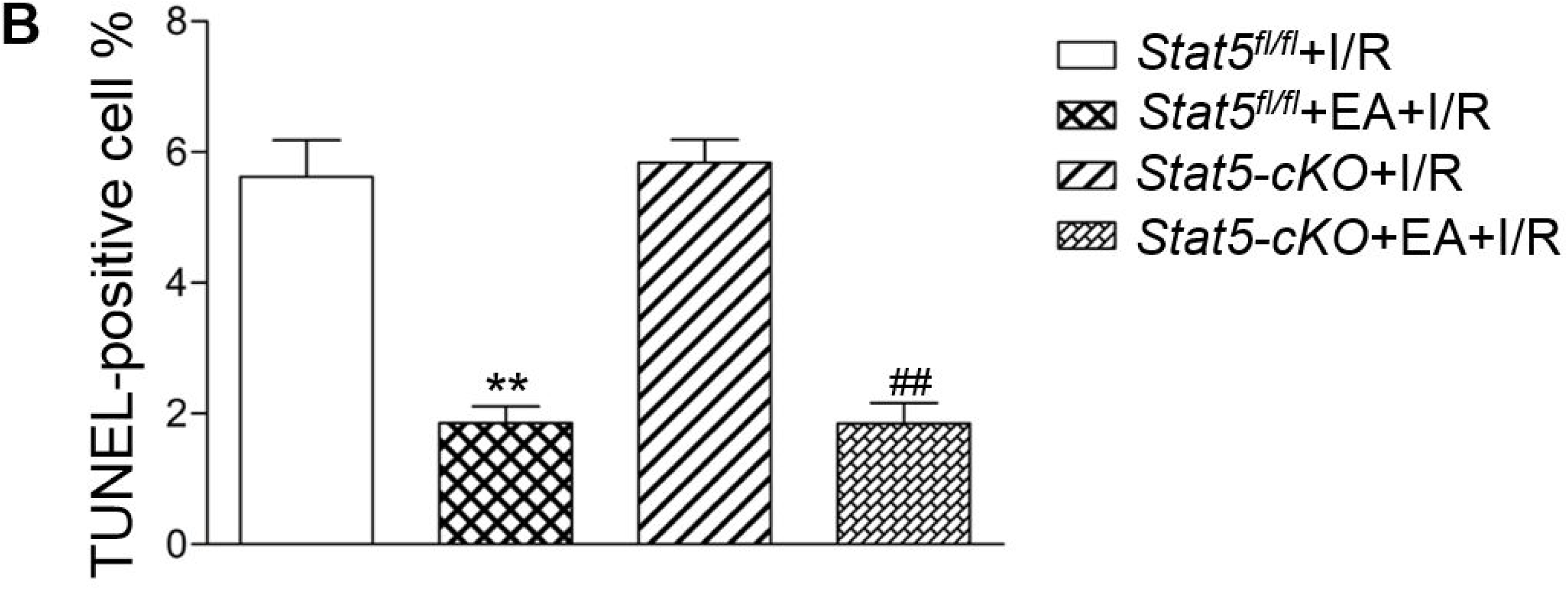
Effects of EAP on apoptosis of myocardial tissues in the *Stat5*^*fl/fl*^ and the *Stat5-cKO* mice. A,B The apoptosis index was measured by TUNEL staining. Data information: Values are mean ± SEM, n = 5. ** *P* < 0.01, compared with the *Stat5*^*fl/fl*^+I/R group, ## *P* < 0.01, in comparison with the *Stat5-cKO*+I/R group.

### 2. EAP activated STAT5 in the *Stat5*^*fl/fl*^ mice but not in the *Stat5-cKO* mice under myocardial I/R condition

To further explore whether the myocardial protection of EAP against I/R injury is STAT5 dependent, we examined protein levels of p-STAT5 in the heart tissues by western blotting. While EAP markedly increased p-STAT5/GAPDH in the *Stat5*^*fl/fl*^ mice compared with the *Stat5*^*fl/fl*^ mice subjected to I/R, EA had no effect on STAT5 activation in the hearts of the *Stat5-cKO* mice (Fig 3A and B). This suggests that STAT5 may participate in the process of EAP protection against myocardial I/R injury.

**Figure 3.**
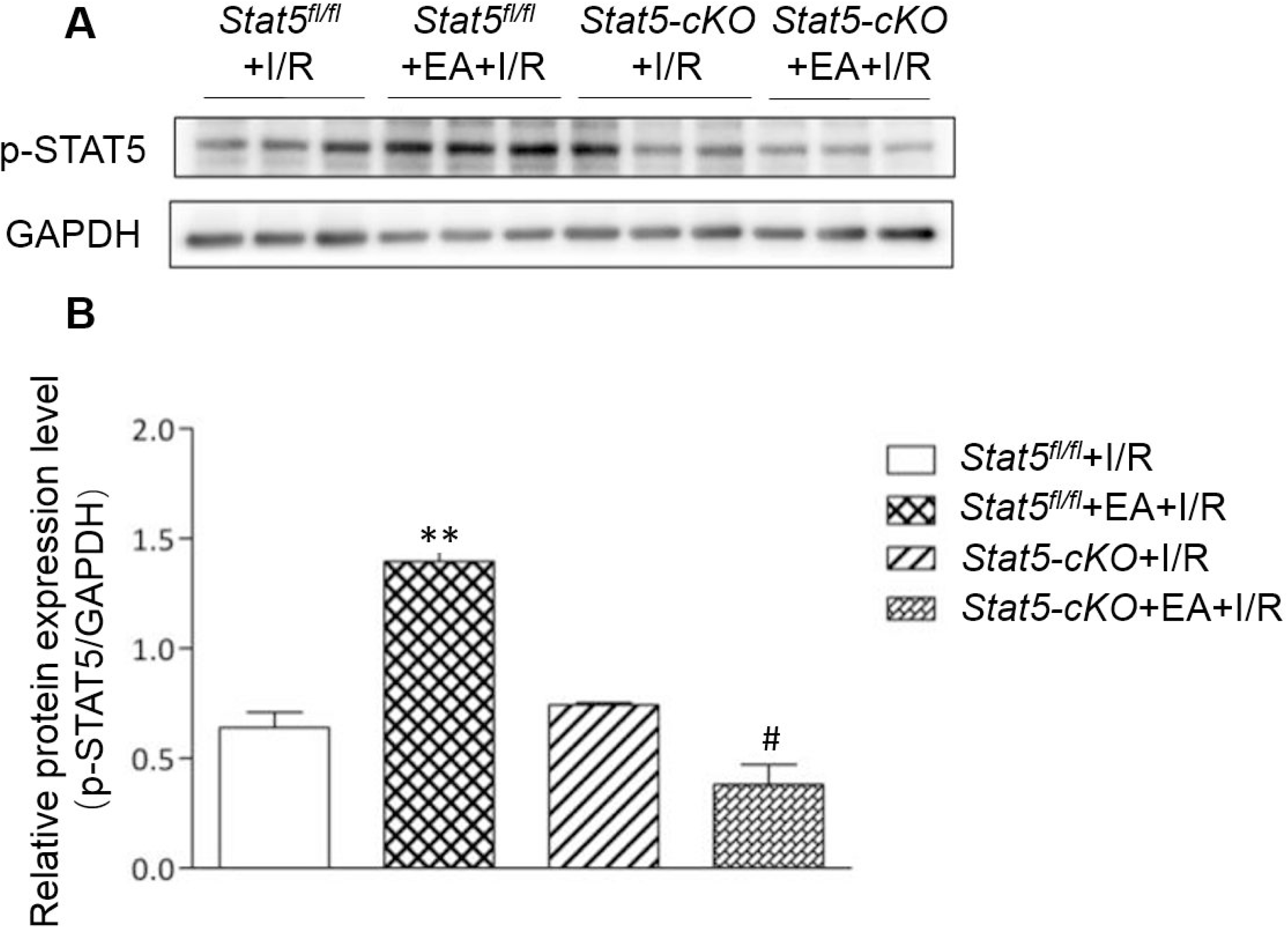
EAP activated STAT5 protein in the heart tissues of the *Stat5*^*fl/fl*^ but not the *Stat5-cKO* mice. A The images of representative western blotting. B Quantitative analysis of p-STAT5 protein in each group. Data information: In B, data are presented as mean ± SEM, data were analyzed by two-way ANOVA, Bonferroni’s multiple comparison test, n=8. ** *P* < 0.01, compared with the *Stat5*^*fl/fl*^+I/R group, # *P* < 0.05, in comparison with the *Stat5-cKO*+I/R group.

### 3. IL-6/gp130/STAT3 signaling was activated by EAP in the absence of *Stat5*

As described in our previous study (Chen *et al*, 2018) and shown in our current observation, STAT5 is a very important molecule in myocardial protection through RIPC or EAP against I/R injury. However, EAP did also display a protective role against myocardial I/R injury when STAT5 was absent in the mouse heart, suggesting that the EAP-induced myocardial protection may be STAT5-independent and some other proteins may be in charge of this function when STAT5 is missing. Considering the possibility of compensation by STAT3, we then evaluated the expression levels of STAT3 protein and p-STAT3 protein in the heart tissues of both *Stat5*^*fl/fl*^ mice and *Stat5-cKO* mice. We discovered that the expression of p-STAT3 was increased in the *Stat5-cKO*+EA+I/R mice compared to the *Stat5-cKO*+I/R group, whereas this was not observed in the *Stat5*^*fl/fl*^ mice (Fig 4A). To understand the mechanism by which STAT3 was activated in this process, we further detected the upstream molecules of STAT3 and found that the mRNA level of IL-6 and gp130 were upregulated in the *Stat5-cKO*+EA+I/R mice compared to those in the *Stat5-cKO*+I/R mice, suggesting that EAP activated the IL-6/gp130/STAT3 pathway in the absence of *Stat5* gene when the heart was exposed to myocardial I/R damage, a process not seen in the presence of *Stat5* (Fig 4B).

**Figure 4.**
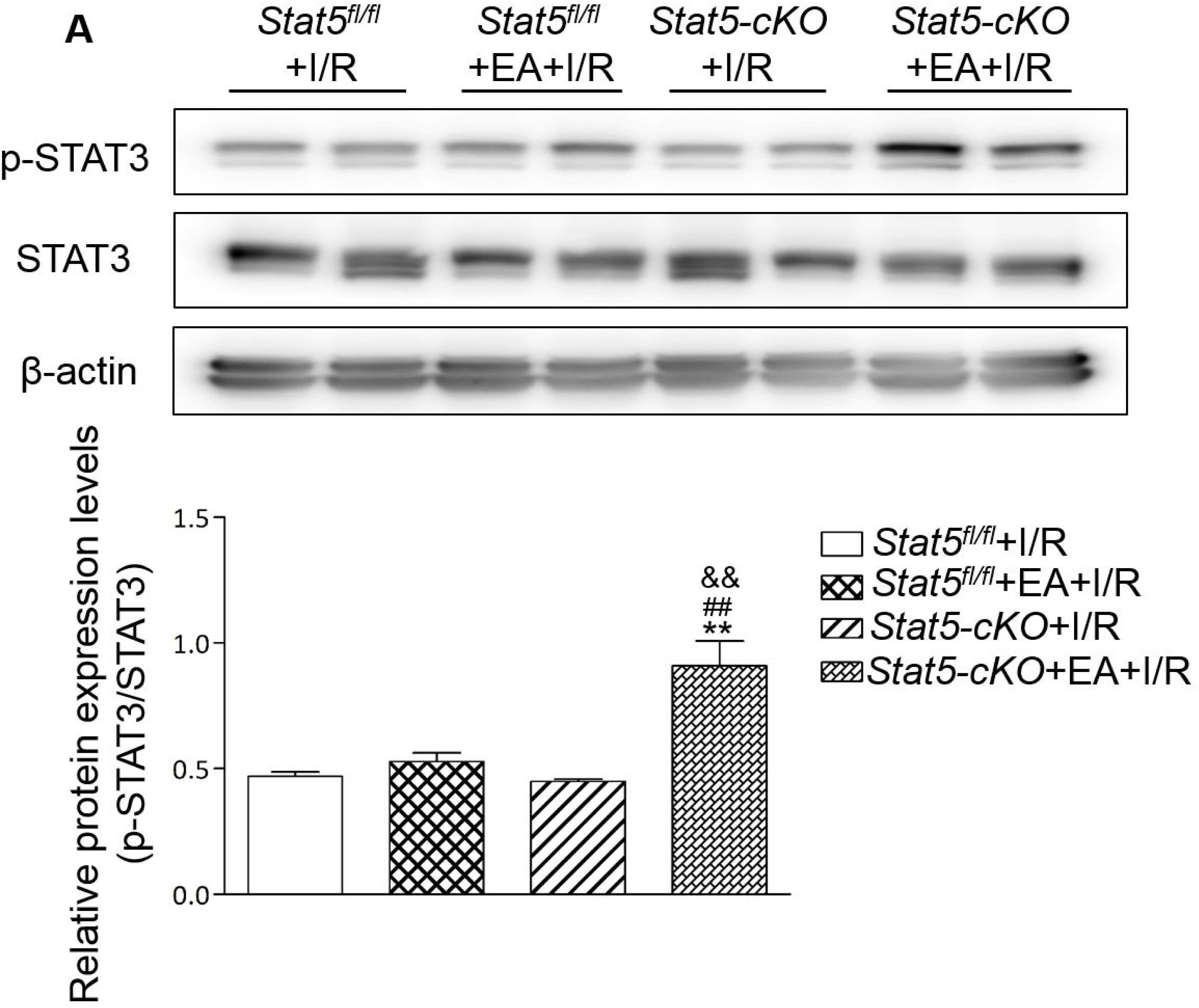
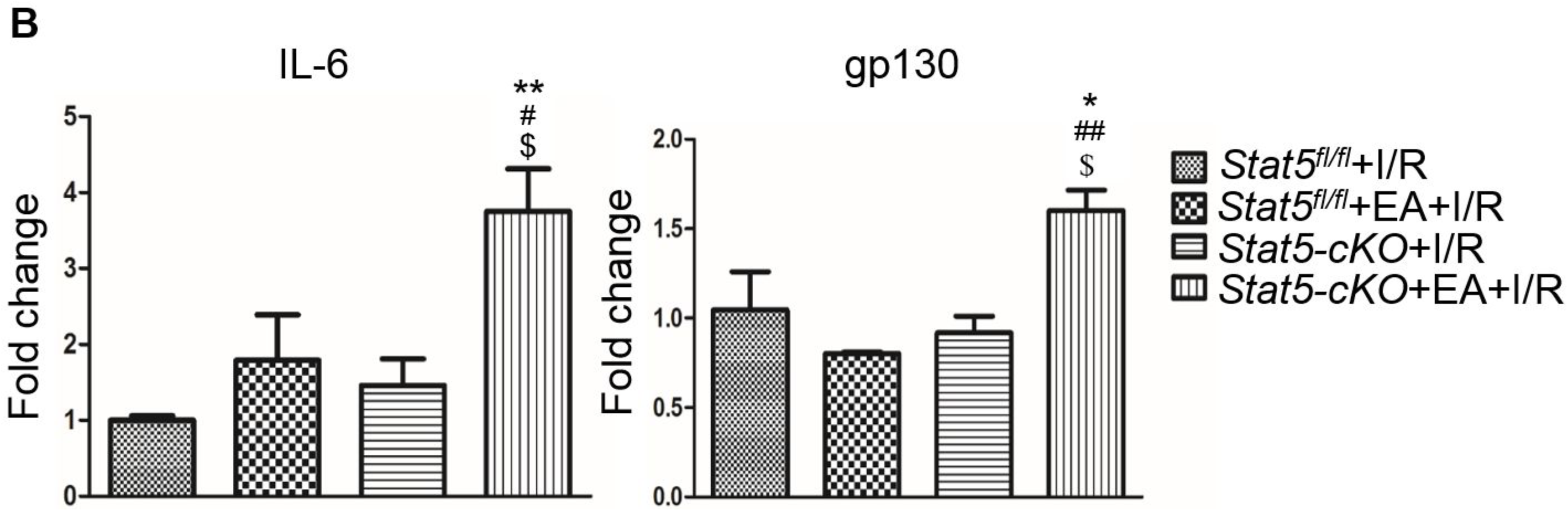
The expression of IL-6/gp130/STAT3 axis related molecules. A The expression of STAT3 and p-STAT3 proteins was detected by western blotting. B The expression of IL-6 and gp130 mRNAs was performed by RT-PCR. Data information: ** *p* < 0.01, compared with the *Stat5*^*fl/fl*^+I/R; ## *P* < 0.01, compared with the *Stat5*^*fl/fl*^+EA+I/R; && *P* < 0.01, compared with the *Stat5-cKO*+I/R, n=4-6. B ** *P* < 0.01, compared with the *Stat5*^*fl/fl*^+I/R group, # *P* < 0.05 compared with the *Stat5*^*fl/fl*^+EA+I/R group, $ *P* < 0.05 compared with the *Stat5-cKO*+I/R group, n=3-4.

### 4. Genome-wide analysis revealed gene expression profiles in both *Stat5* ^*fl/fl*^ mice and *Stat5-cKO* mice with or without EAP followed by myocardial I/R injury

To find the candidate genes participating in the EAP-induced protection against myocardial I/R injury, RNAs were extracted from the heart tissues and RNA-seq was performed using next generation high-throughput sequencing. With the Cufflinks package, we filtered out the top 30 differentially expressed genes (DGEs) by fold changes through comparing the *Stat5*^*fl/fl*^+I/R group with the *Stat5*^*fl/fl*^+EA+I/R group and the *Stat5-cKO*+I/R group with the *Stat5-cKO*+EA+I/R group (Table 1A and B). First, we compared DEGs between the *Stat5*^*fl/fl*^+I/R group and the *Stat5*^*fl/fl*^+EA+I/R group and between the *Stat5*-*cKO*+I/R group and the *Stat5-cKO*+EA+I/R group. Venn diagrams were drew based on the list of filtered DEGs among four groups (Fig 5). The results showed that, compared with the *Stat5*^*fl/fl*^+I/R group, 1052 genes were differentially expressed in the *Stat5*^*fl/fl*^+EA+I/R group, and 1039 DEGs were obtained by comparing the *Stat5*-*cKO*+I/R group and the *Stat5-cKO*+EA+I/R group, in which 133 genes were overlapped by these two clusters. Among these 4 groups, only two genes, Hspa1a and Pttg1 were found to belong to all four groups, suggesting that STAT5-dependent genes and EAP-regulated genes were located in different categories. We further analyzed these genes and further tried to understand mechanisms by which myocardial I/R pathology occurs in the presence or absence of *Stat5* and how EAP protects myocardial I/R injury through regulating gene expressions. We found that among the EAP up-regulated genes and down-regulated genes in the presence of STAT5 (*Stat5*^*fl/fl*^+I/R vs *Stat5*^*fl/fl*^+EA+I/R), lots of genes (such as Fosb, Fos, Cxcl5, Cxcl, Egr1, Egr2, Nr4a3, Socs3, Ccn5, Myl4, Zhx2, Dkk3, and Dynll1) have been reported to participate in the protection against the myocardium ischemia, I/R, cardiac hypertrophy or hypoxia in the literatures (Wu *et al*, 2017; Kubota *et al*, 2019; Wang *et al*, 2018; Sun *et al*, 2019; Ma *et al*, 2020; Oba *et al*, 2012; Tian *et al*, 2020; Stobdan *et al*, 2015; Bos *et al*, 2012; Zhai *et al*, 2018; El-Magd *et al*, 2017). These DEGs belong to many functional pathways, such as the JAK/STAT signaling pathway, the TNF signaling pathway, apoptosis, or the NF-kappa B signaling pathway. In the other hand, when STAT5 is absent, we found that among the top 30 DEGs generated by comparing the *Stat5-cKO+*EA*+*I/R group with the *Stat5-cKO+*I/R group (Table 1B), Rps6, Mmp3, pttg1, Rac2 had a strong relationship with the IL-6/STAT3 signaling pathway, as reported previously in the literatures (Dern *et al*, 2019; Wu *et al*, 2020; Calamaras *et al*, 2015; Sharma *et al*, 2019; Abilleira *et al*, 2006; Zhu & Sun *et al*, 2017; Huang *et al*, 2018; Lai *et al*, 2017; Shirakawa *et al*, 2018; Wen *et al*, 2015).

**Table 1.**
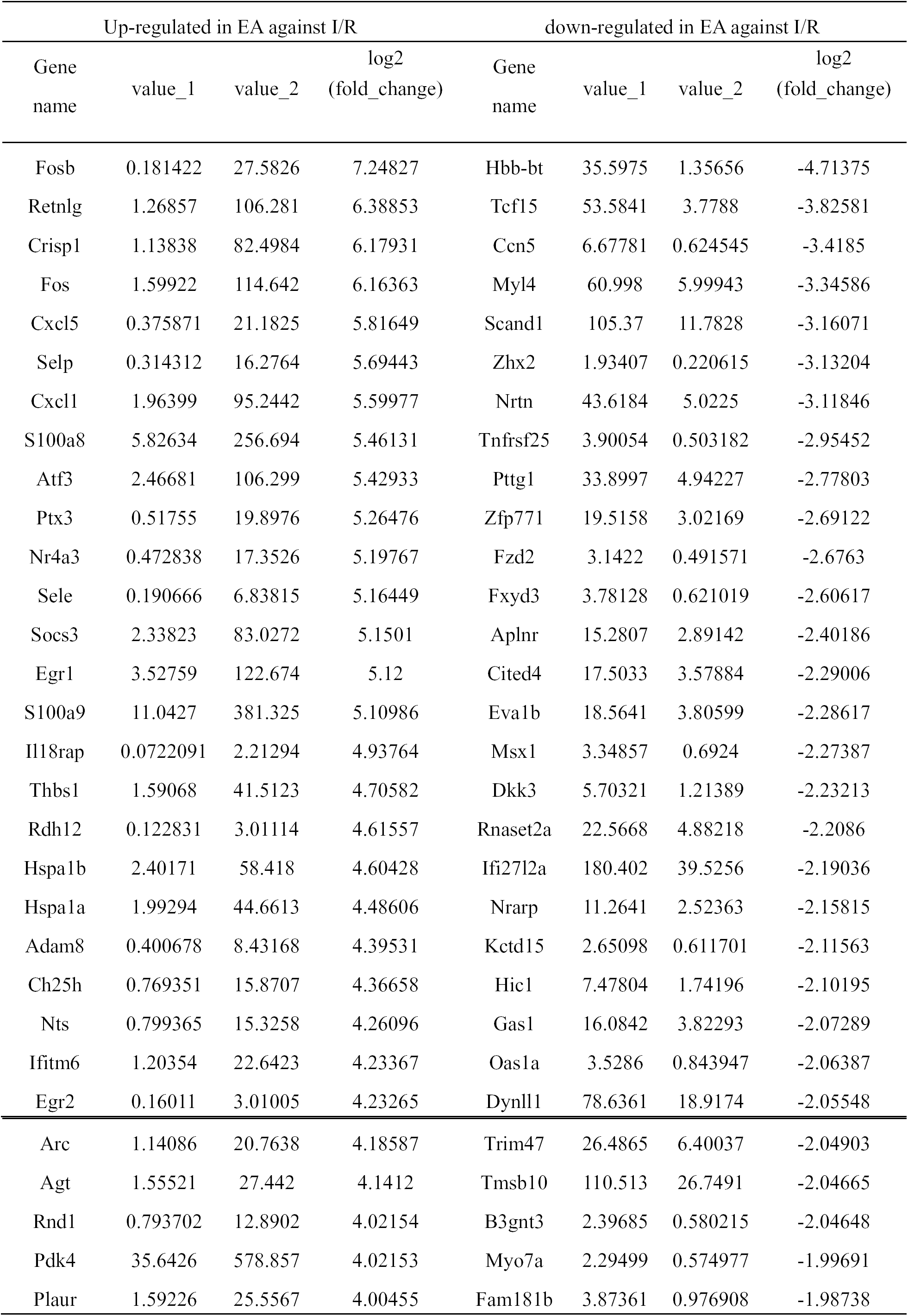
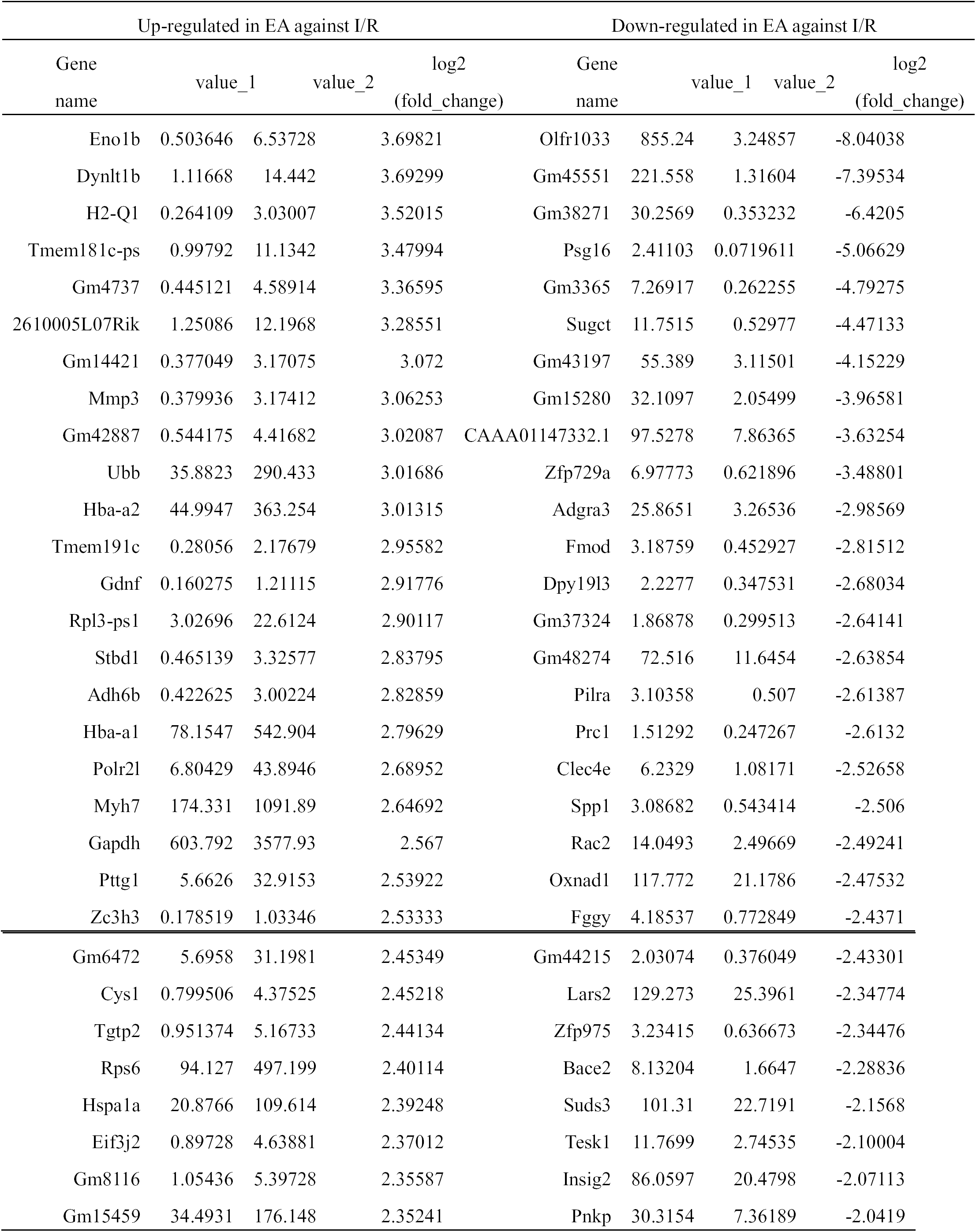
The top 30 differentially expressed genes with a log2 (FC) > |±1|and q value<0.05 A. The top 30 differentially expressed genes obtained from comparing *Stat5*^*fl/fl*^*+*EA+I/R vs B. The top 30 differentially expressed genes obtained from comparing *Stat5-cKO*+EA+I/R vs. *Stat5-cKO*+I/R

**Figure 5.**
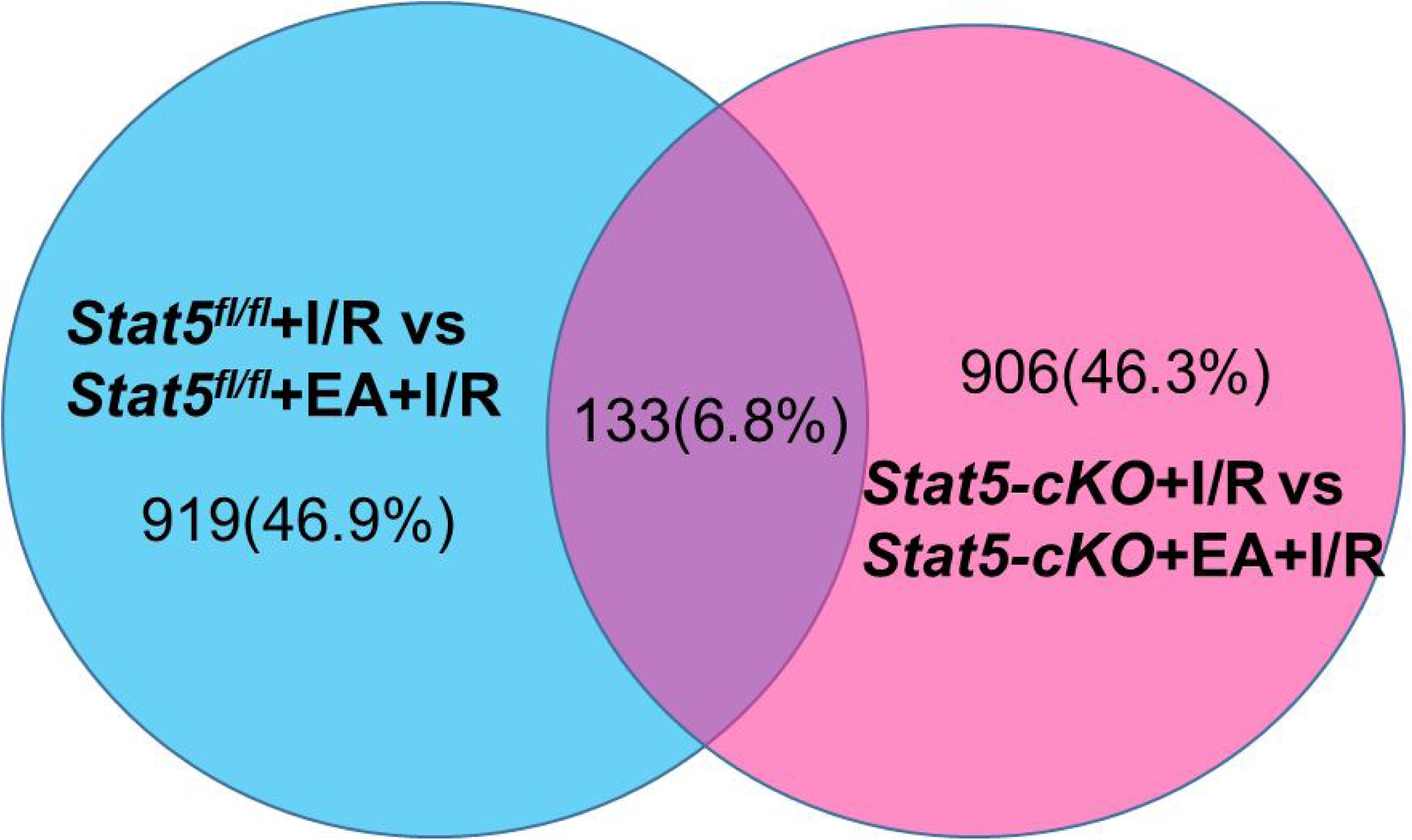
Venn diagrams and clustering analysis of RNA-seq results. Data information: Venn diagrams were drawn based on our RNA-seq data sets. Blue circle indicates the numbers of genes up- and down-regulated in the *Stat5*^*fl/fl*^*+*EA+I/R group (vs. the *Stat5*^*fl/fl*^+I/R group); pink circle represents the numbers of up- or down-regulated genes in the *Stat5-cKO*+EA+I/R group (vs. *Stat5-cKO*+I/R group). 133 genes were overlapped by these two clusters.

To further understand the potential pathways involved in the STAT5-related DEGs and the EAP-modified DEGs under I/R, we then carried out pathway analyses for these DEGs. Encyclopedia of Genes and Genomes (KEGG) pathway analyses were performed using DAVID Bioinformatics Resources, and the top 20 pathways were outlined in Fig 6.

**Figure 6.**
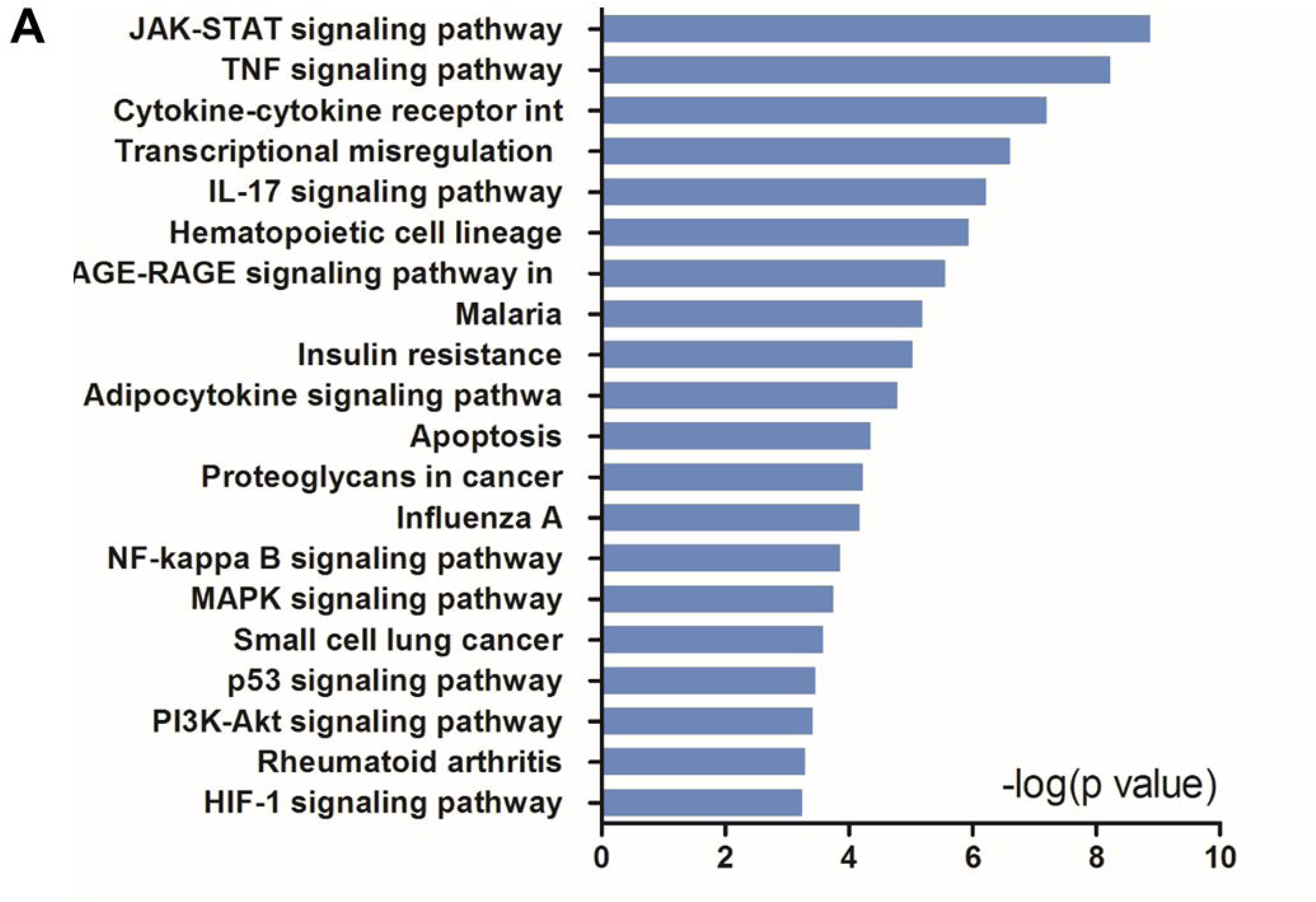
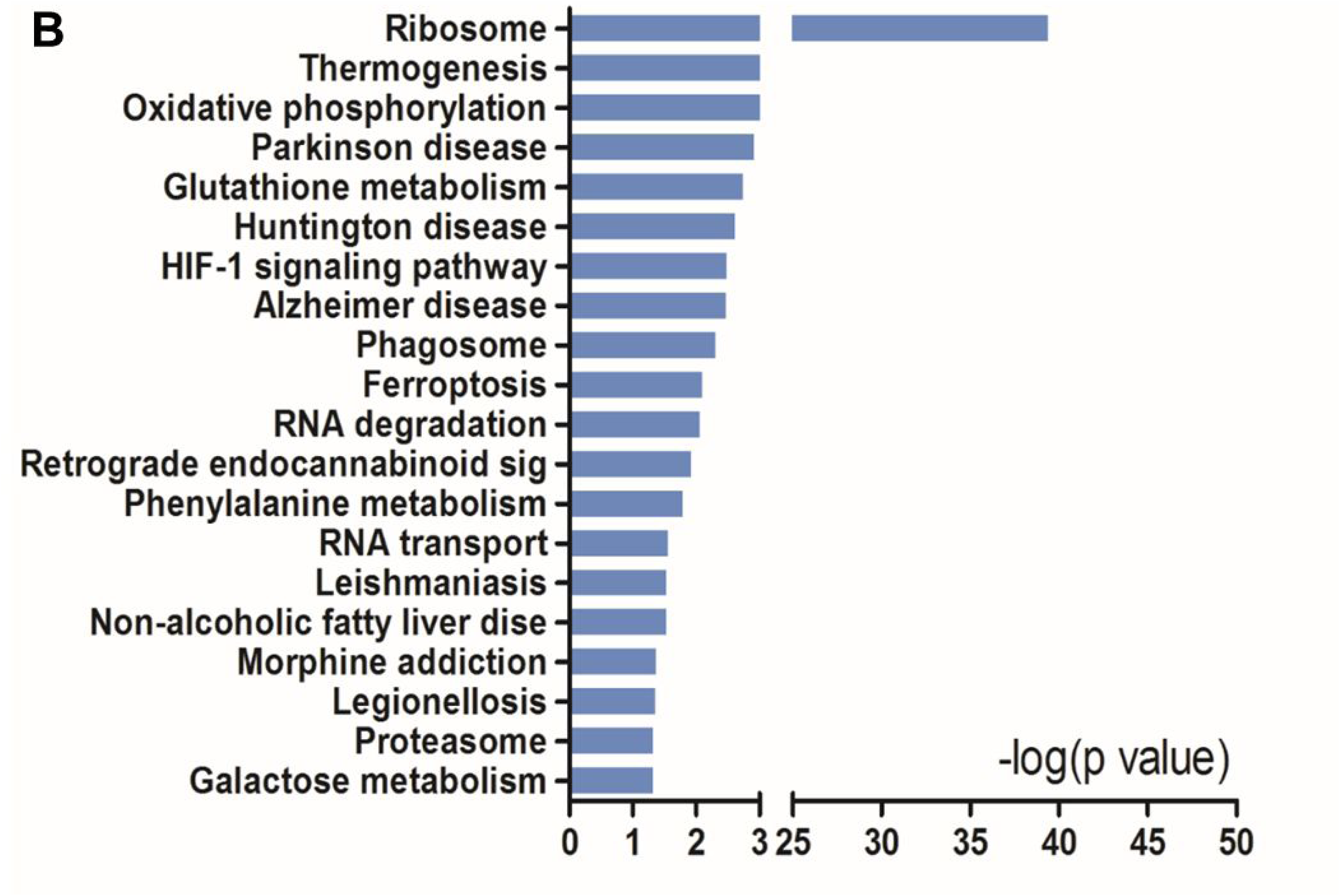
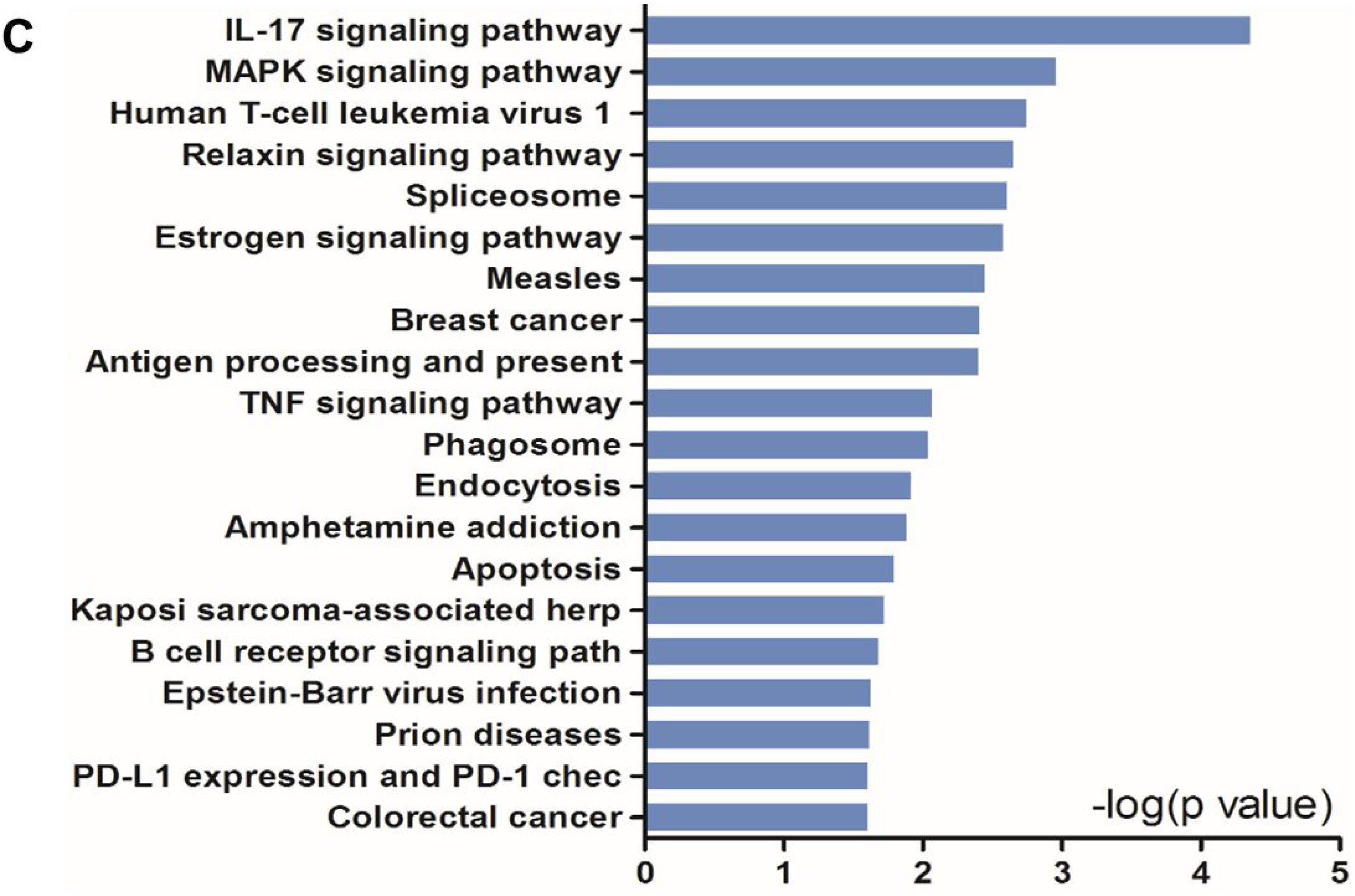
KEGG pathway analysis of up- and down-regulated genes in the heart tissues in the presence or absence of *Stat5* (A–C). A The top 20 KEGG pathways were drawn from the 919 DEGs genes by comparing the *Stat5*^*fl/fl*^+I/R group and the *Stat5*^*fl/fl*^+EA+I/R group shown in Figure 5. B The top 20 KEGG pathways were drawn from the 906 DEGs genes by comparing *Stat5-cKO*+I/R group and *Stat5*-*cKO*+EA+I/R group shown in Figure 5. C The top 20 KEGG pathways were drawn from the 133 co-regulated genes shown in Figure 5.

KEGG pathway analyses suggested that, in the presence of *Stat5*, EAP-activated genes were mainly enriched in the JAK-STAT signaling pathway, the TNF signaling pathway, cytokine-cytokine receptor interaction, the IL-17 signaling pathway, the NF-kappa B signaling pathway, and the MAPK signaling pathway (Fig 6A). They may work together to protect the myocardium from I/R injury (Castejon *et al*, 2019; Yuan *et al*, 2019; Świerkot *et al*, 2015). In contrast, in the *Stat5-cKO* mice, the myocardial protection of EAP mainly concentrated in Ribosome pathways, thermogenesis, and the oxidative phosphorylation pathway (Fig 6B). Moreover, we also analyzed the top 20 KEGG pathways of the 133 overlapped genes (Fig 6C), showing that the EAP-regulated part of the STAT5-independent genes were mainly located in inflammation related pathways (such as the IL-7 signaling pathway, human T-cell leukemia virus 1 infection, antigen processing and presentation, and the TNF signaling pathway).

### 5. Apoptotic and survival signaling were regulated by EAP only in the presence of STAT5

Based on the above genome-wide profiling data, we detected that EAP could activate apoptotic and survival signaling in mice with I/R injury. To further validate these findings, we studied apoptotic and survival related protein expression in myocardial tissues of the *Stat5*^*fl/fl*^ and the *Stat5-cKO* mice with EAP. The results showed that the expression of Bcl-2 and Bcl-xL were significantly increased in the *Stat5*^*fl/fl*^+EA+I/R group compared with the *Stat5*^*fl/fl*^+I/R group (*P*<0.05), whereas the expression of Cyt C did not change between these two groups (Fig 7). In contrast, we did not detect significant differences in these protein expressions in the hearts of *Stat5-cKO* mice, with or without EAP, suggesting that STAT5 is required for regulating anti-apoptotic signaling induced by the EAP stimulus. We then detected the level of IL-10, an important cytokine in cardio-protection, and its related proteins, PI3K, AKT, and p-AKT (Fig 8). The results showed that IL-10 and p-AKT were elevated by EAP in the presence but not absence of STAT5, however IL-10 was up-regulated in the hearts of both *Stat5*^*fl/fl*^ mice and *Stat5-cKO* mice with EAP. These results indicate that the protective effect of EAP against myocardial I/R injury in survival signaling is partially STAT5-dependent.

**Figure 7.**
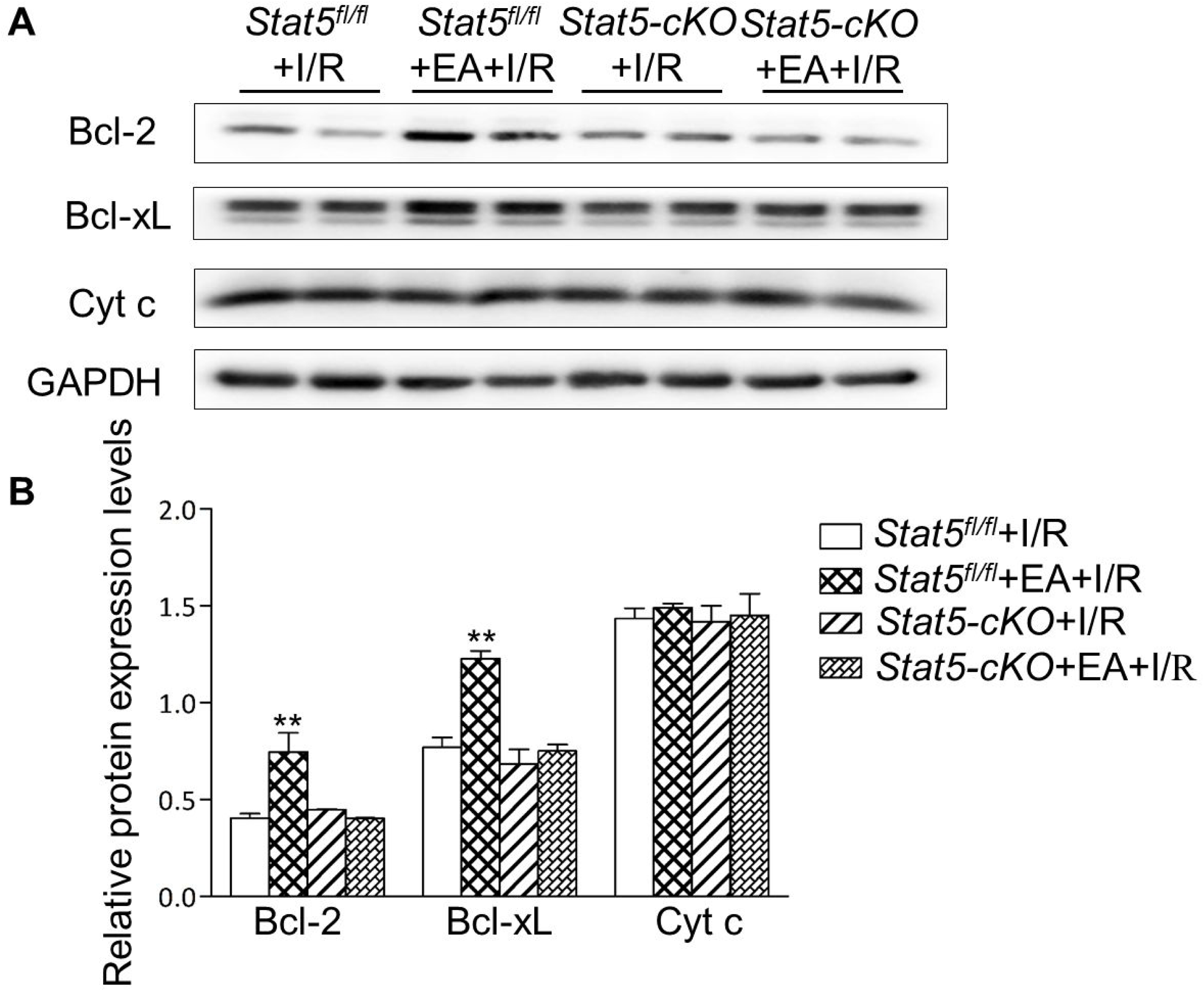
The expression of apoptosis-related proteins. A,B Western blotting was used to detect the level of Bcl-2, Bcl-XL and Cyt C in each group. Data information: ** *P* < 0.01, compared with *Stat5*^*fl/fl*^+I/R group, n=6.

**Figure 8.**
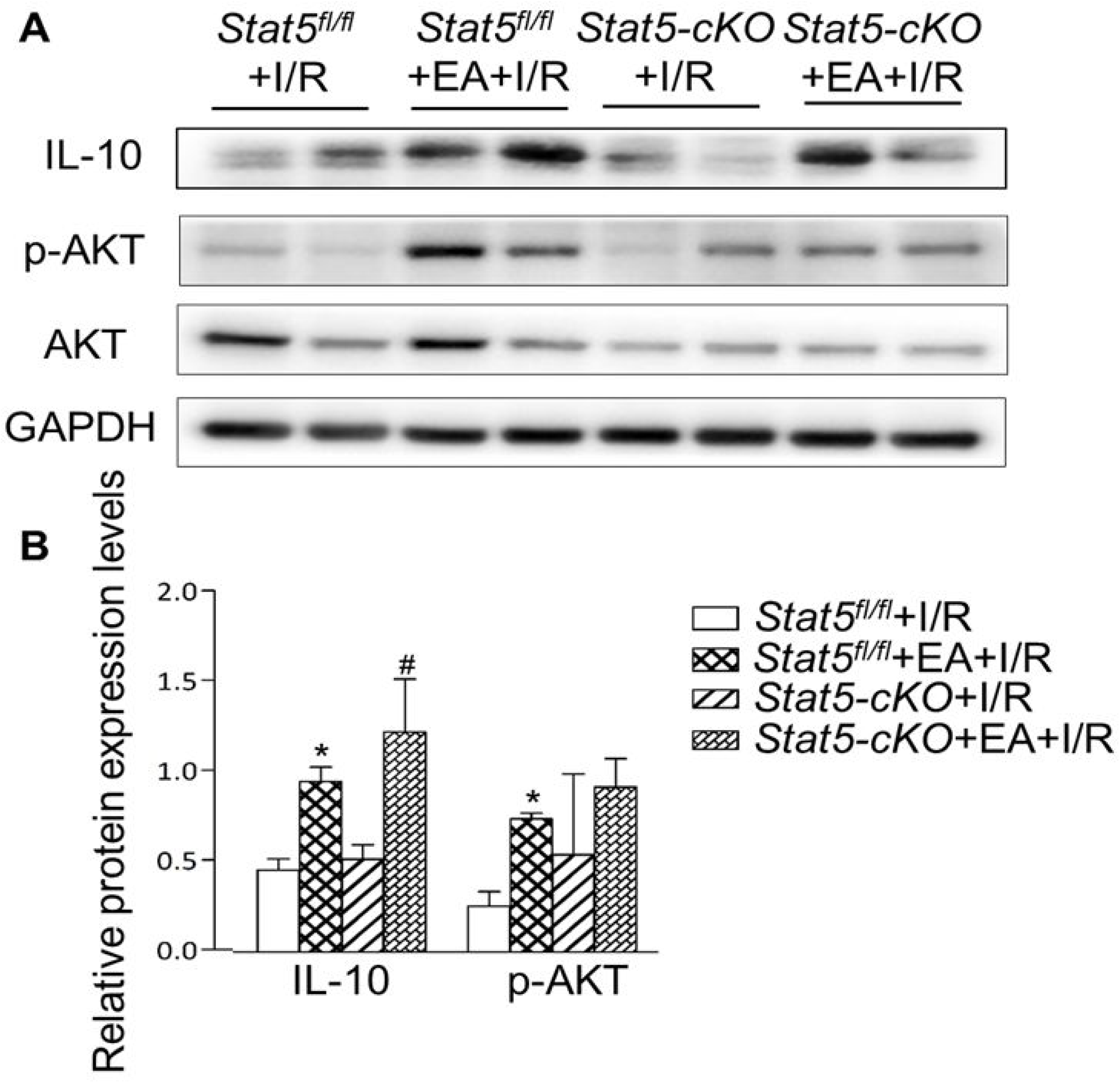
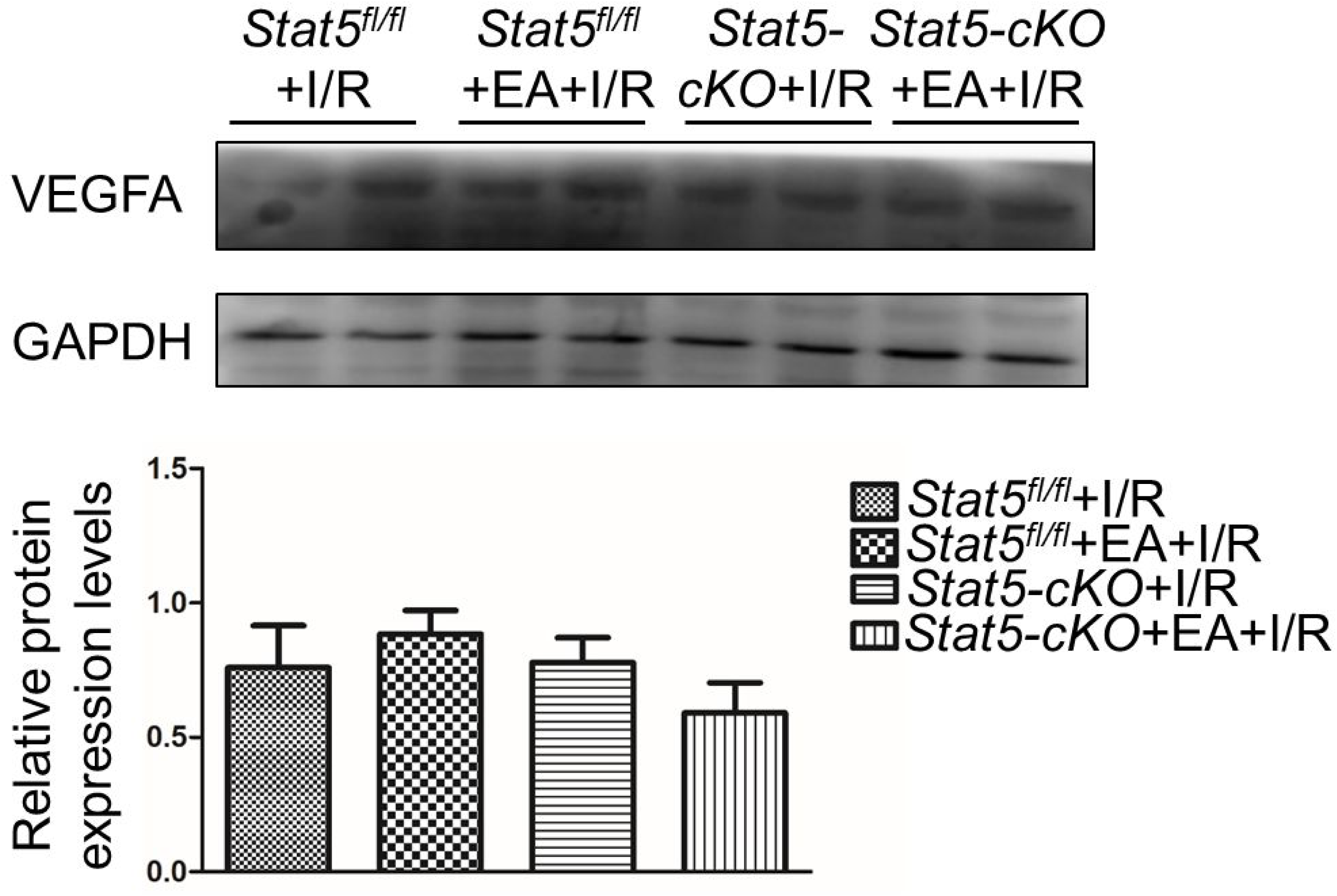
The expression of survival signaling-related proteins. A,B Western blotting was used to detect the level of IL-10, p-AKT, and AKT in each group. Data information: * *P* < 0.05, compared with *Stat5*^*fl/fl*^+I/R group, # *P* < 0.05, in comparison with *Stat5-cKO*+I/R group, n=6.

## Discussion

Ischemic heart disease is still the major cause of premature mortality and disability worldwide (Gupta & Wood, 2015). As a clinically effective method against myocardial I/R injury, early coronary reperfusion reduces the infarct size, but reperfusion by revascularization initiates a cascade of events that can accelerate and extend post-ischemic injury (Binder *et al*, 2015; Yellon & Hausenloy, 2017). Remote ischemia preconditioning (RIPC) has been confirmed to be an effective and clinically applicable perioperative method to reduce the risk of myocardial injury (Hausenloy *et al*, 2019; Ekeloef *et al*, 2019). In our previous study, we have found that STAT5 plays a key role in the RIPC-mediated late cardio-protection through anti-apoptotic signaling and the PI3K/AKT survival pathway (Chen *et al*, 2018). Similar to RIPC, EA pretreatment at acupoint PC6 like a late stimulating can also protect myocardium under certain disease conditions by stimulating multiple functional pathways.

In the present study, we explored the role of STAT5 in EAP-induced myocardial protection against ischemia-reperfusion by employing cardiomyocyte-specific *Stat5-cKO* mice. We observed that EAP could reduce the infarct size and myocardial cell apoptosis in both *Stat5*^*fl/fl*^ and *Stat5-cKO* mice (Fig 1 and Fig 2), suggesting that STAT5 is not required in the cardioprotection of EAP against myocardial I/R injury. To understand how EAP played a protective role in the loss of *Stat5*, we performed RNA sequencing with the I/R injured heart tissue. We found that EAP can increase the mRNA expressions of Fosb, Fos, Cxcl5, Cxcl1, Egr1, Egr2, Nr4a3, and Socs3, which were reported to be involved in anti-apoptosis, anti-inflammation, antioxidation, and STAT3/5 signaling when STAT5 is intact. Fosb and Fos, as two components of AP-1, can protect against myocardial I/R injury via anti-apoptosis, anti-oxidative stress, and anti-inflammatory activation (Wingelhofer *et al*, 2018; Walker *et al*, 2014; Walker *et al*, 2013). Cxcl5 and Cxcl1 are two members in the CXC family of chemokines and have been shown to improve cell survival after myocardial injury through the promotion of wound healing by changing neutrophil infiltration and activating the phosphatidylinositol 3-kinase pathway (Cai *et al*, 2020; Zhou *et al*, 2020). Egr1 and Egr2 belong to early growth response protein 1/2. They can induce myocardial injury by regulating myocardial autophagy, cell death, and the progression of myocardial fibrosis (Wang *et al*, 2018; Cao *et al*, 2010). Nr4a3, a member of the NR4A orphan nuclear receptor, plays a protective role during acute myocardial infarction by suppressing inflammatory responses via the JAK2-STAT3/NF-kB pathway (Obana *et al*, 2010; Walker *et al*, 2014). Socs3, a negative effector of STAT3 signaling, is an NF-kappaB/IKK-induced gene. IKK-mediated NF-kappaB activation can coordinately illicit negative effects on STAT signaling (Gopinath, 2017). Aside from the above up-regulated genes, Hspa1a and Hspa1b are also important factors during the restoration of myocardial I/R injury (Jiang *et al*, 2014; Wilhide *et al*, 2011) and appear to be up-regulated by EAP. In addition, among the down-regulated DEGs, Ccn5 is an anti-hypertrophic and anti-fibrotic factor during adverse cardiac remodeling. It can be regulated by activating JAK/AKT/STAT3-signaling in luminal-type (ER-positive) breast cancer (BC) cells (Haque *et al*, 2018). Myl4 (myosin light polypeptide 4) is a key gene for atrial contractile, electrical, and structural integrity (Udoko *et al*, 2016). Zhx2, as one of the transcription factors, can improve macrophage survival and pro-inflammatory functions in atherosclerotic lesions (Tian *et al*, 2020; Alfonso-Jaume *et al*, 2006). Hic1, a transcriptional repressor that modulates the expression of cell-cycle genes, can affect heart function and be regulated by the IL-6/STAT3 pathway (Wen *et al*, 2019; Sulston *et al*, 2017). Gas1, pttg1, Nrtn, and Tnfrsf25 have also been identified as key molecules in the heart tissue by regulating tissue formation, metabolism, apoptosis of cardiomyocytes, and the interaction of cytokine-cytokine receptor in the JAK/STAT pathway (Tang *et al*, 2019; Jiang *et al*, 2019; Saddic *et al*, 2018).

In short, many of the genes and pathways found in our RNA-seq have been attributed to MI or I/R injury and are regulated by EAP in the *Stat5*^*fl/fl*^ mice. This indicates that EAP can mimic RIPC and play a protective role against I/R injury by altering functional gene expression in the presence of STAT5.

Intriguingly, in the *Stat5-cKO* mice, EAP regulated different DEGs, of which Rps6, Mmp3, Pttg1, and Rac2 are involved in the IL-6/STAT3 signaling pathway. In addition, ribosomal, thermogenesis, and oxidative phosphorylation pathways are activated by EAP in the absence of STAT5. Among these DGEs, Mmp3, known as matrix metallopeptidase 3, encodes a member of the matrix metalloproteinase family of the extracellular matrix-degrading enzymes that are involved in tissue remodeling, wound repair, progression of atherosclerosis, and tumor invasion (Abilleira *et al*, 2006). Recent findings indicate that the binding of STAT3 to the MMP promoter promotes the transcription of Mmp3 gene, which accounts for IL-6-induced MMP gene activation (Zhu & Sun, 2017). Pituitary tumor transforming 1 (pttg1) has been reported as an oncogene that is originally cloned from rat pituitary tumor cells. Huang *et al*. have demonstrated that pttg1 expression is regulated by IL-6 via the direct binding of activated STAT3 to the pttg1 promoter in LNCa P cells. Rac2, one of the Rac family members, is expressed mainly in the hematopoietic cells (Huang *et al*, 2018). Rac protein can promote glioblastoma tumor sphere-induced angiogenesis in the zebrafish exnotransplantation model. Knockdown of Rac protein reduces the tumorigenesis in the mouse model *in vivo* (Lai *et al*, 2017). Lai *et al*. have detected reduced STAT3 activation in the Rac down-regulated glioblastoma cells without affecting STAT5 activation. Osteopontin (OPN) is also known as secreted phosphoprotein 1 (Spp1). High levels of intracellular galectin-3 expression are essential for transcriptional activation of Spp1 in STAT3-mediated polarization toward M2 macrophages after MI (Shirakawa *et al*, 2018; Wen *et al*, 2015). The phosphorylation sites of ribosomal protein S6 (Rps6) have been mapped to five clustered residues, which play an important role in protein synthesis in cardiac myocytes and cardiac function (Dern *et al*, 2019; Wu *et al*, 2020; Calamaras *et al*, 2015; Sharma *et al*, 2019). Through RNA-seq profiling, we expect different mechanisms were involved in EAP protection against myocardial I/R injury between *Stat5*^*fl/fl*^ and *Stat5-cKO* mice. Combined with the molecular biological data in this study, these results support our hypotheses that EAP may activate STAT3 in the absence of STAT5 and function as a protective approach for I/R.

In fact, multiple studies have demonstrated that in the absence of a given STAT member, receptors will recruit other STAT members instead (Hennighausen *et al*, 2018; Yu *et al*, 2010; Valle & Soto *et al*, 2020; Hosui *et al*, 2009; Hin *et al*, 2020; Friedbichler *et al*, 2012). STAT5 and STAT3 are two proteins of STAT family that show high homology in their functional domains, can be activated by different mechanisms, and bind to distinct loci to regulate specific target gene expression (Wingelhofer *et al*, 2018). STAT3 and STAT5 proteins can also bind to the same regulatory oncogenic loci, resulting in compensatory or antagonistic signaling (Walker & Xiang, 2014; Walker *et al*, 2013). Nevertheless, the role of STAT5 and STAT3 in myocardial I/R injury by EAP have not been studied yet. Interestingly in our study, p-STAT3 protein has significantly increased in the *Stat5-cKO*+EA+I/R group compared to the *Stat5*^*fl/fl*^+EA+I/R group (Fig 4), suggesting that EAP activates STAT3, which then contributes to protecting the myocardium against I/R injury in the *Stat5-cKO* mice. Furthermore, RNA-seq data suggest that the ribosome pathway is significantly activated by EAP. It may also link to the cardioprotection in the absence of *Stat5*. In this pathway, we found that Rps6 and Rpl3-ps1 are upregulated by EAP in the *Stat5-cKO* mice. They were reported before to be regulated by the IL-6/STAT3 signaling (Dern *et al*, 2020; Meyuhas *et al*, 2015). To confirm this, we determined the mRNA expression in the IL-6/gp130 receptor system, which is an important activator of STAT3, and have found that the mRNA expressions of gp130 and IL-6 are increased by EAP only in the *Stat5-cKO* mice (Fig 4B), suggesting that when STAT5 is deleted, IL-6/gp130/STAT3 signaling gets activated to play role in the protection against myocardial I/R.

Growing evidence has demonstrated the favorable and protective role of STAT3 in the heart (Harhous *et al*, 2019; Nakao *et al*, 2020). To understand the mechnisms by which STAT3 contributes to protection of EAP against I/R imjury, we determined the apoptotic and survival signaling in the heart tissue of the mice. STAT3 is involved in decreasing cardiac I/R injury by reducing apoptosis or increasing anti-apoptotic signaling, increasing expression of cardioprotective proteins, decreasing ROS generation, and inhibiting autophagy (Harhous *et al*, 2019). We found that EAP promoted the expression of anti-apoptotic proteins Bcl-2 and Bcl-xl and p-AKT in the *Stat5*^*fl/fl*^+I/R mice, but not in the *Stat5-cKO+*I/R mice. However, the expression of IL-10 protein was increased in both the *Stat5*^*fl/fl*^ and the *Stat5-cKO* mice when EAP was applied followed by I/R injury. IL-10 is one of the important anti-inflammatory cytokines which can be produced by most cells and affect the growth and differentiation of various hematopoietic cells and increase cell proliferation, angiogenesis, and immune evasion (Zhen *et al*, 2018; Hodge *et al*, 2005). Our previous study has shown that IL-10 RIPC can activate the expression of IL-10, p-AKT, Bcl-2, and Bcl-xl to protect the myocardium (Chen *et al*, 2018). Recently, Takahashi J *et al*. has shown that interleukin-22, one member of the IL-10 cytokine family, activates myocardial STAT3 signaling pathway and prevents myocardial I/R injury in the mouse model of ischemia reperfusion injury (Takahashi *et al*, 2020). Other studies have also shown that the IL-6 and IL-10 family of cytokines are the main mediators that activate intrinsic JAK/STAT3 signaling to induce the transcription of genes enabling survival and proliferation of cells (Harhous *et al*, 2019; Huynh *et al*, 2017). Activated STAT3 can regulate the expression of genes (such as MMP2, MMP9, and Ubc13) and therefore underpin the molecular cross-talk between these genes (Pipicz *et al*, 2018). When the *Stat5-cKO* mice were given a stress of myocardial I/R injury in this study, Mmp3, Ubb, and Myh7 genes that are closely correlated with STAT3 pathway were altered by EAP (Table 1b), therefore STAT3 might have played a vital role in cardio-protection by controlling these gene’s expression. In addition, some studies have identified that activation of STAT3 can improve the expression VEGF (Huynh *et al*, 2019; Johnson *et al*, 2018). We have detected the expression of VEGF-A, but there is no difference among our four groups (Appendix Fig 9). To a certain extent, these results show that the activation of STAT3 by EAP may be not enough for elevating angiogenesis in the *Stat5-cKO* mice.

In summary, the present study demonstrates that EAP approach to protecting against myocardial I/R injury by reducing the myocardial infarct area and activating anti-apoptotic and survival signaling. STAT5 is involved in this process but the protection is not STAT5 dependent. STAT3 may compensate the function of STAT5 by activating the IL-6/gp130/STAT3 signaling pathway in the absence of STAT5. This study suggests, for the first time, that EAP can mimic RIPC but function more effectively in cardio-protection against I/R injury through multiple pathways.

## Materials and Methods

### Antibodies and reagents

Antibodies for p-STAT5 (Try694), p-STAT3 (Try705), STAT3, p-AKT (Ser473), AKT, Cytochrome c (Cyt c), Bcl-xL, Bcl-2, and GAPDH were purchased from Cell Signaling Technology. Antibodies for IL-10, and β-actin were purchased from Abcam (Cambridge, UK). The in-situ cell-death detection kit to assess apoptosis, POD (TUNEL), was purchased from Roche (Lewes, UK). The triphenyltetrazolium chloride (TTC) was purchased from Sigma-Aldrich (St. Louis, MO, USA).

### Conditional and inducible cardiomyocyte-specific *Stat5-cKO* mice

As previously described, the *Stat5* mice (*Stat5*^*fl/fl*^) were kindly provided by Dr. Hennighausen (NIDDK, NIH), and Tnnt2-Cre male mice (*Tnnt2*^*Cre*^) were obtained from Bin Zhou (Shanghai Institutes for Biological Sciences of the Chinese Academy of Sciences) as gifts (Chen *et al*, 2018). The *Stat5* knockout mice (*Stat5-cKO*) were generated by mating these two genotypes. Doxycycline hyclate (Sigma-Aldrich, St. Louis, MO, USA) was administrated through adding into the drinking water of mice at a concentration of 2 mg/ml for 7 days. The method used for genotyping has been described in our previous article (Chen *et al*, 2018).

### Study groups

The mice were divided into four groups: *Stat5*^*fl/fl*^+I/R, *Stat5*^*fl/fl*^+EA+I/R, *Stat5-cKO+*I/R, and *Stat5-cKO*+EA+I/R. *Stat5*^*fl/fl*^+I/R and *Stat5-cKO+*I/R groups were subjected to left anterior descending (LAD) coronary artery occlusion for 30min, and then reperfused for 180 min. *Stat5*^*fl/fl*^+EA+I/R and *Stat5-cKO*+EA+I/R groups were subjected to EAP 7 days before the LAD ligation. All murine studies were carried out in accordance with the EU Directive 2010/63/EU for the protection of animals used for experimental purpose. All experiments were approved by the Institute for Animal Care and Use Committee at Nanjing University of Chinese Medicine.

### In vivo experiments

Prior to the myocardial I/R injury experiment, *Stat5*^*fl/fl*^+EA+I/R and *Stat5-cKO*+EA+I/R mice were pretreated with EA for a total of 7 days. EA was performed at billateral PC6 (also called Neiguan) acupoints based on the previous study (Huang *et al*, 2014). The PC6 acupoint is located in the interosseal muscles between the radius and ulna of the forelimb, 3 mm proximal to the wrist crease, according to the textbook of experimental acupuncture in animals (Huang *et al*, 2014). Mice were anaesthetized with isoflurane (5%) and maintained with 1-2% isoflurane in pure oxygen. The sterilized disposable stainlesssteel acupuncture needles (0.18mm×13mm, Beijing Zhongyan Taihe Medical Instruments Factory, Beijing, China) were inserted a depth of 1-2mm into the muscle layer at bilateral PC6 simultaneously using Han’s EA instrument (Han Acuten, WQ1002F, Beijing, China) and alternating dense and disperse mode, with a frequency of 2/15 Hz at an intensity level of 0.5-1mA, at a stimulation period of 20 minutes, once a day, for a total 7 days. The mice in the *Stat5*^*fl/fl*^+I/R group and the *Stat5-cKO*+I/R group were restrained for 20 minutes without EA stimulation.

The I/R operation was performed as described in the previous studies (Chen *et al*, 2018; Huang *et al*, 2014). Briefly, all the mice were given 5% isoflurane and then maintained with 2% isoflurane in a mixture of 70% N2O and 30% O2 for anesthesia. Under the anesthetized state, the mice were subjected to a left thoracotomy and LAD ligation with a slipknot using 6-0 silk sutures transiently at 2 to 3 mm below the left auricle, resulting in arterial occlusion, as evidenced by myocardial blanching and electrocardiographic abnormalities (ST-segment elevation and QRS complex widening) (Chen *et al*, 2018). After 30 min, reperfusion was performed by quickly releasing and removing the suture and continued for 3h. In the sham-operation group, the same procedure was performed except for the LAD ligation. Mice were sacrificed after surgery and the heart specimens were harvested.

### Determination of infarct size

At the end of the protocol, sections of mouse heart were perfused for 1-2 min with 2ml of 2% TTC (Sigma-Aldrich Co., St. Louis, MO, USA) in phosphate-buffered saline (PBS) and then incubated in an identical solution at 37°C for 15 min. After incubating, slices were placed in 4% (v/v) paraformaldehyde at 4°C for 12h. TTC stained area (red, normal area) and non-stained areas (white, infarct area) were photographed. The infarcted area was quantified using Image-Pro Plus 6.0 software (NIH, USA). All analyses of infarct size were performed by two investigators who were blinded with the information regarding group assignments.

### Apoptosis measurements

TUNEL staining method was used to detect cell apoptosis of cardiac tissue in each group. All the protocols were the same as previously described (Chen *et al*, 2018). Heart tissues were harvested and embedded with OCT (Thermo Scientific^™^, USA). The 8μm thick tissues were subjected to TUNEL staining according to the manufacturer’s instructions for an In-Situ Cell Death Detection Kit (Cat. 11684817910, Roche Diagnostics, Indianapolis, IN, USA). Sections were then visualized with a fluorescence microscope (Nikon, Japan) with parallel positive control (DNase-I) and negative control (label solution only).

### Western blotting

Whole ventricle samples were lysed with RIPA buffer supplemented with a protease inhibitor cocktail and a phosphatase inhibitor cocktail (Chen *et al*, 2018). Homogenates were centrifuged at 14,000×g for 10 min at 4 °C, and supernatants were stored at -80 °C until further uses. Protein concentrations were determined using a BCA protein assay (Pierce). Protein was mixed with 5× Laemmli loading buffer and heated at 95 °C for 10 min. Equal amount of protein was subjected to SDS-PAGE and transferred to polyvinylidene fluoride (PVDF) membranes. Immunoblots were performed using appropriate primary antibodies against Bcl-xL (1:1000, Cell Signaling, #2762), Bcl-2 (1:1000, Cell Signaling, #3498), Cyt c (1:1000, Cell Signaling, #6808), Phospho-STAT5 (1:1000, Cell Signaling, #9662), STAT5(1:1000, Cell Signaling, #3421), Phospho-STAT3 (1:1000, Cell Signaling, #2775), STAT3 (1:1000, Cell Signaling, #2612), Phospho-AKT (1:1000, Cell Signaling, #9662), AKT (1:1000, Cell Signaling, #9608), IL-10 (1:1000, Cell Signaling, #4351), β-actin (1:1000, Cell Signaling, #1721), or GAPDH (1:1000, Cell Signaling, #1752) overnight at 4°C, then incubated with a secondary antibody for two hours at room temperature. Western blot band intensities were quantified using SuperSignal West Pico Chemiluminescent substrate (Pierce).

### q-PCR analysis

Three micrograms of RNA were converted to cDNA using reverse transcriptase and random primers (11121ES60, Yeasen Biotech Co., Ltd., China). For PCR analysis, the samples were amplified in duplication using SYBR Green (Vazyme, Q431-02) with 200 nM of gene-specific primers and run on the CFX amplifier (MX3000P, Stratagene, La Jolla, CA, USA) according to the manufacturer’s protocol. Data were analyzed by the threshold cycle (Ct) relative-quantification method. The primer sequences were as follow: interleukin-6 (IL-6, forward: GACTTCACAGAG GATACCACCC, reverse: GACTTCACAGAGGATACCACCC); gp130 (forward: GAGCTTCGAGCCATCCGGGC, reverse: AAGTTCGAGCCGCGCTGGAC); Actin (forward: GGTGAAGACGCCAGTAGAC, reverse: TGCTGGAAGGTGGACAGTGA).

### RNA-seq and computational analysis for RNA-seq data

To further explore possible changes in transcriptome profile caused by EAP against myocardial I/R injury in the *Stat5* knockout mice, RNA-seq for the mouse heart tissues were performed using next generation high throughput sequencing method (Illumina HiSeq 2000, Illumina). Total RNAs were isolated. The RNA-seq protocol was described in our previous study (Fu *et al*, 2017).

Data analysis was performed as previously described (Fu *et al*, 2017). Quality of raw sequencing data was assessed by FASTQ files. We used the Cufflinks program and the Cuffdiff program to assemble individual transcripts and differential transcript expression analysis. The pathways were analyzed using the DAVID Bioinformatics Resources. Genes with lower than 1.0 FPKM (average fragments perkilobase of transcript per million fragments mapped) filtered out. Up-regulated and down-regulated genes were defined as a relative transcription level of above Log2 fold change (FC) ≥| ±1| and *P* value <0.05.

### The paper explained

#### Problem

Our recent study has shown that signal transducer and activator of transcription 5 (STAT5) plays a critical role in RIPC, and RIPC mediates cardioprotection by activating anti-apoptotic and cardiomyocyte-survival signaling in a STAT5-dependent manner (Chen *et al*, 2018). Studies, including our previous work, have shown that using electro-acupuncture before the event of ischemia-reperfusion, which is also called electro-acupuncture pretreatment (EAP), could protect cardiomyocytes by reducing the myocardial infarct size and regulating some molecular signaling (Huang *et al*, 2014; Lu *et al*, 2016). Given that the main molecule involved in RIPC in human patients is STAT5 (Cheung *et al*, 2006; Chen *et al*, 2018), we question whether EAP shares the same mechanism with RIPC and plays a preconditioning-like role as RIPC does on I/R injury.

#### Result

The present study demonstrates that EAP can protect against myocardial I/R injury by reducing the myocardial infarct area and activating anti-apoptotic and survival signaling. STAT5 is involved in this process but the protection is not STAT5 dependent. STAT3 may compensate the function of STAT5 by activating the IL-6/gp130/STAT3 signaling pathway in the absence of STAT5. We revealed, for the first time, that EAP can mimic RIPC but function more effectively in cardio-protection against I/R injury through multiple pathways.

#### Impact

This study will provide new scientific experimental data for acupuncture protecting against I/R injury.

#### Statistics

SPSS18.0 statistical software was utilized for statistical analysis. All data were presented as the mean ± standard error of mean (SEM). The significance of the differences was determined by Student’s t-test or one-way analysis of variance (ANOVA) with the least significant difference post hoc test when equal variances were assumed or with Bonferroni test post hoc when equal variances were not assumed. *P* < 0.05 was considered statistically significant.

## Data availability

The data supporting this study is available from the corresponding author upon reasonable request. RNA-Seq raw data is avilable in the National Center for Biotechnology Information (NCBI) Gene Expression Omnibus (GEO) with the accession number GSExxxxx (https://www.ncbi.nlm.nih.gov/geo/query/acc.cgi?acc=GSExxxxx).

Expanded View for this article is available online

## Acknowledgements

This research was supported by the National Key R&D Program of China (No. 2019YFC1709003), the National Natural Science Foundation of China (Grant No. 81870224). We thank Dr. Wanxin Liu (Washington DC, USA) for language editing.

## Author contribution

Bing-mei Zhu and Xin-yue Jing conceived and supervised experiments. Hui-hui Guo, Xin-yue Jing, Hui Chen and Bing-mei Zhu wrote and edited the manuscript. Hui Chen and Hui-hui Guo performed experiments and analyzed data. Hou-xi Xu carried out the bioinformatic analyses for RNA-seq.

## supplementary material

manuscriptID_Figure 9.

